# Relationship of drought severity index with climatic factors on the Tibetan Plateau

**DOI:** 10.1101/494815

**Authors:** Xiangtao Wang, Guangyu Zhang, Jingsheng Wang

## Abstract

It remains unclear how drought is varying under climatic change. Quantifying the relationship between drought and climatic factors is crucial for predicting future drought risk under global climatic change. Correlations of annual drought severity index (DSI) with climatic factors were examined from 2000 to 2011 on the Tibetan Plateau. Spatially averaged DSI increased with increasing precipitation and minimum relative humidity, but decreased with increasing sunshine. The degrees of correlation between DSI and climatic factors varied with vegetation types. The change magnitude of DSI decreased with increasing temperature, precipitation and vapor pressure, but increased with increasing wind speed and sunshine. Therefore, clarifying the correlation between drought and climatic change need consider ecosystem types and their local climate on the Tibetan Plateau.

## 1 Introduction

Drought is an important climatic event and is generally linked to water availability, temperature, sunshine and wind speed [1-4]. Generally, water availability plays an important role for terrestrial ecosystems and drought has an adverse effect on vegetation growth [2,5-7]. Several studies have focused on how drought is changing under climatic change; however, non-uniform results on this question have been reported based on models and observations [3,8,9]. Knowledge of the relationship between drought and climatic change is crucial for predicting future changes in aridity and managing drought severity [2,3,10].

The Tibetan Plateau is one region most sensitive to climatic change [11-13]. Approximately two-third of the Tibetan Plateau are arid and semiarid regions [5]. The arid and semiarid regions may get drier under climatic change [14]. Under the background of global warming, the Tibetan Plateau is experiencing obvious warming and the warming magnitude is much greater than the global average [11,12]. It is expected that climatic warming could exacerbate drought when drought occurs [3]. Several recent studies indicate that decreased magnitude of soil moisture caused by experimental warming is negatively related to warming magnitude of soil temperature [10,15,16]. These findings imply that the drought risk may be more extensive and its magnitude may be greater on the Tibetan Plateau compared to other regions of the world. Actually, [17] have found that the increasing drought risk on the Tibetan Plateau is much higher than the average in China.

Climatic change varies with vegetation types on the Tibetan Plateau [18-20]. To our knowledge, no studies have focused on the comparing the correlations between inter-annual variation of drought index and climatic change among vegetation types over the Tibetan Plateau. Moreover, previous studies have mainly discussed the correlation of drought index with temperature and precipitation rather than other climatic factors (e.g. vapor pressure, relative humidity, wind speed and sunshine hours) on the Tibetan Plateau [21,22]. Recently, the Moderate Resolution Imaging Spectroradiometer (MODIS) developed a new drought index, called drought severity index (DSI), using the MODIS-derived actual and potential evapotranspiration, and normalized difference vegetation index [2], which facilitates analyses of the drying conditions and associated relationships with climatic factors at global and regional scales. In this study, we used MODIS DSI data and meteorological data to: (1) analyze and compare the inter-annual variation of DSI and its relationships with climatic factors among the main terrestrial ecosystems; and (2) analyze whether climatic warming has a negative effect on DSI on the Tibetan Plateau.

## 2 Materials and methods

### 2.1 Study area

The Tibetan Plateau is called ‘Third Pole of the Earth’ and its unique features includes high altitude (mean altitude above 4000 m), high air transparency, strong solar radiation, thin air and low temperature [23,24]. Alpine meadows, alpine steppes, temperate steppes, forests, shrubland and croplands are main vegetation types [15,25].

### 2.2 MODIS DSI data

In this study, DSI data were obtained from the Moderate Resolution Imaging Spectroradiometer (MODIS) drought severity index product. The spatial and temporal resolutions are 0.05°×0.05° and one year, respectively. The images in 2000–2011 on the Tibetan Plateau were adopted. Inconsistent with most previous drought indices, the vegetation responses are incorporated in the DSI algorithm [2]. The MODIS global DSI product has been validated [2].

### 2.3 Climatic data

We obtained climatic data from China Meteorological Data Sharing Service System. In this study, 69 meteorological stations (22 in alpine meadows, 8 in alpine steppes, 4 in temperate steppes, 7 in forests, 12 in shrublands and 16 in croplands) were used. Climatic data included five temperature factors [(annual average air temperature (Ta), maximum air temperature (MAT), minimum air temperature (MIT), extreme maximum air temperature (EMAT), and extreme minimum air temperature (EMIT)], two precipitation factors [annual total precipitation (TP), and maximum precipitation (MAP)], four humidity factors [annual average vapor pressure (Ea), relative humidity (RH), minimum relative humidity (MIRH), and vapor pressure deficit (VPD)], two sunshine factors [annual percentage of sunshine (SP) and total sunshine hours (SH)] and annual average wind speed (WS).

### 2.4 Data analysis

The vegetation type map (1:1000000 scale) was firstly transformed into a raster file (0.05°×0.05° spatial resolution) before any other associated analyses. We used a trend analysis to analyze changes in DSI, Ta, MAT, MIT, EMAT, EMIT, TP, MAP, Ea, RH, MIRH, VPD, SP, SH and WS. We used a correlation analysis to analyze the relationship between DSI and climatic factors [18]. All the spatial analyses were performed using the ArcGIS (version 9.3) [18].

## 3 Results and discussion

### 3.1 Climatic factor changes

Annual climate became warming and drying from 2000 to 2011 based on these meteorological records on the Tibetan Plateau (Figure S1). In detail, based on the data from all the 69 meteorological stations, spatially averaged Ta, MAT and MIT increased at 0.07, 0.09 and 0.07 °C·yr^−1^, respectively. Spatially averaged MAP exhibited an increase at 0.33 mm·yr^−1^. Spatially averaged Ea and RH exhibited a decrease at −0.04 hpa·yr^−1^ and −0.01 yr^−1^, respectively. Spatially averaged VPD increased at 0.01 hpa·yr^−1^.

Climate changes varied with vegetation types (Figure S1), which was in line with previous studies [18,19,26]. In general, the climate showed a warming and drying from 2000 to 2011 across meteorological stations in alpine meadows, alpine steppes, forests and shrublands, but a drying in temperate steppes and croplands. The climate also became warming if considering the significant increase of MAT or MIT in croplands and temperate steppes. A dimming occurred in alpine steppes and temperate steppes, but a brightening occurred in shrublands.

In detail, spatially averaged Ta increased at 0.08, 0.09, 0.07 and 0.08 °C·yr^−1^ in alpine meadows, alpine steppes, forests and shrublands, respectively. Spatially averaged MAT increased at 0.09, 0.07, 0.11 and 0.11 °C·yr^−1^ in alpine meadows, croplands, forests and shrublands, respectively. Spatially averaged MIT increased at 0.09, 0.12, 0.07, 0.07 and 0.07 °C·yr^−1^ in alpine meadows, alpine steppes, temperate steppes, forests and shrublands, respectively. Spatially averaged TP increased at 5.95 and 6.50 mm·yr^−1^ in alpine steppes and temperate steppes, respectively. Spatially averaged MAP increased at 0.38 and 0.72 mm·yr^−1^ in alpine meadows and shrublands, respectively. Spatially averaged Ea decreased at −0.02, −0.03, −0.07 and −0.07 hpa·yr^−1^ in alpine steppes, croplands, forests and shrublands, respectively. Spatially averaged RH decreased at −0.006, −0.007, −0.003, −0.004, −0.007 and −0.009 yr^−1^ in alpine meadows, alpine steppes, temperate steppes, croplands, forests and shrublands, respectively. Spatially averaged MIRH increased at 0.003 yr^−1^ in temperate steppes and at 0.002 yr^−1^in croplands. Spatially averaged VPD increased at 0.07, 0.08, 0.04, 0.07, 0.12 and 0.13 hpa·yr^−1^ in alpine meadows, alpine steppes, temperate steppes, croplands, forests and shrublands, respectively. Spatially averaged SP increased at 0.002 yr^−1^ in shrublands, but decreased at −0.003 yr^−1^ in temperate steppes. Spatially averaged SH increased at 8.49 hours·yr^−1^ in shrublands, whereas spatially averaged SH decreased at −12.1 and −11.8 hour·yr^−1^ in alpine steppes and temperate steppes, respectively. Spatially averaged WS increased at 0.02 m·s^−1^·yr^−1^ in croplands.

### 3.2 DSI changes

Spatially averaged DSI change during the past 12 years varied with vegetation types (Figure 1). Only spatially averaged DSI in forests decreased at approximately –0.039 yr^−1^. This was in line with the fact that the decreasing magnitude of TP and the increasing magnitude of VPD in forests was the largest among the six vegetation types.

Inter-annual DSI change during the past 12 years varied among the 69 stations (Figure 2). Eight stations exhibited a decrease at rates from −0.13 to −0.09 yr^−1^. In contrast, only the Wudaoliang station exhibited an increase at 0.10 yr^−1^. This finding was in line with previous studies which indicated that the change of water availability varied with stations during the past years on the Tibetan Plateau [18,19].

The Southern Tibetan Plateau exhibited a drying trend, while the Northern Tibetan Plateau exhibited a wetting trend (Figure 2). This finding was in line with previous studies [5,18,27].

The DSI change decreased with increasing Ta, MAT, MIT, EMAT, EMIT, TP, MAP and Ea, but increased with increasing SP, SH and WS (Figure S3). These findings implied that DSI was more responsive to warming in colder environments, to water availability in drier environments and to sunshine conditions in lighter environments. The DSI was also more responsive when the wind speed is high. Therefore, clarifying the correlation between DSI and water availability need consider the local climate conditions.

The DSI change exhibited a positive relationship with TP change, but negative correlations with changes of MAT, EMAT, EMIT, SP and SH (Figure 3). Likewise, recent studies found that a higher experimental warming resulted in a greater environmental drying [10,15,16]. These findings indicated that warming magnitude and change magnitude of water availability exhibited quite the opposite effects on DSI change. The negative effect of sunshine conditions change magnitude on DSI change also dampened the positive effect of water availability change magnitude on DSI change. These findings suggested that DSI change was correlated with the changes and background values of climatic factors.

Climatic factor changes and background values differed among the six vegetation types or among the 69 meteorological stations (Figure S1, S2), which was in line with previous studies conducted on the Tibetan Plateau [18-20]. Therefore, different inter-annual DSI variations may be attributed to different climatic changes and background values of climatic factors among the six vegetation types or among the 69 meteorological stations.

### 3.3 Relationships between DSI and climatic factors

Over all the 69 meteorological stations, spatially averaged DSI was positively related to spatially averaged TP and MIRH, but negatively correlated to SP and SH (Table 1). Therefore, DSI can mirror the condition of water availability and the increase of sunshine probably resulted in drying over the meteorological stations [2].

**Table 1.** Correlation coefficients for spatially averaged annual drought severe index (DSI) with spatially averaged annual average temperature (Ta), maximum temperature (MAT), minimum temperature (MIT), extreme maximum temperature (EMAT), extreme minimum temperature (EMIT), total precipitation (TP), maximum precipitation (MAP), average vapor pressure (Ea), average relative humidity (RH), minimum relative humidity (MIRH), average vapor pressure deficit (VPD), percentage of sunshine (SP), sunshine hours (SH) and wind speed (WS) from 2000 to 2011 at 69 meteorological stations on the Tibetan Plateau

| Vegetation<br>Types | Ta | MAT | MIT | EMAT | EMIT | TP | MAP | Ea | RH | MIRH | VPD | SP | SH | WS |
| --- | --- | --- | --- | --- | --- | --- | --- | --- | --- | --- | --- | --- | --- | --- |
| Alpine meadows | -0.11 | -0.42 | 0.26 | -0.43 | -0.08 | 0.59* | 0.20 | 0.67* | 0.50 | 0.71** | -0.35 | -0.85*** | -0.87*** | 0.49 |
| Alpine steppes | 0.28 | -0.07 | 0.58* | -0.36 | 0.20 | 0.70* | 0.19 | 0.07 | -0.17 | -0.35 | 0.23 | -0.74** | -0.74** | 0.19 |
| Temperate steppes | 0.20 | 0.10 | 0.25 | 0.04 | 0.57 | 0.42 | -0.19 | 0.29 | -0.16 | 0.07 | 0.34 | -0.23 | -0.19 | 0.18 |
| Croplands | -0.34 | -0.48 | -0.14 | -0.55 | -0.05 | 0.43 | -0.22 | 0.36 | 0.50 | 0.26 | -0.49 | -0.90*** | -0.91*** | 0.26 |
| Forests | -0.71** | -0.77** | -0.73** | -0.63* | -0.69* | 0.54 | 0.24 | 0.74** | 0.78** | -0.04 | -0.76** | -0.66* | -0.65* | -0.02 |
| Shrublands | -0.66* | -0.78** | -0.51 | -0.61* | -0.36 | 0.66* | -0.43 | 0.69* | 0.75** | 0.83*** | -0.73** | -0.75** | -0.75** | 0.03 |
| All types | -0.28 | -0.49 | -0.03 | -0.40 | -0.11 | 0.61* | -0.18 | 0.54 | 0.50 | 0.74** | -0.42 | -0.86*** | -0.86*** | 0.31 |

\*, \*\* and \*\*\* indicate $p < 0.05$ , $p < 0.01$ and $p < 0.001$ , respectively.

Spatially averaged DSI increased with increasing TP in alpine meadows, alpine steppes and shrublands (Table 1). Spatially averaged DSI exhibited a positive correlation with Ea in alpine meadows, forests and shrublands (Table 1). Spatially averaged DSI decreased with increasing VPD, but increased with increasing RH in forests and shrublands (Table 1). Spatially averaged DSI exhibited a positive relationship with MIRH in alpine meadows and shrublands (Table 1). These findings implied that DSI can mirror the dynamic of water availability at ecosystem scale. However, the degree of correlations between DSI and climatic factors related to water availability varied with vegetation types (Table 1). Therefore, clarifying the correlation between DSI and water availability need consider ecosystem types.

Spatially averaged DSI decreased with increasing Ta, MAT and EMAT in forests and shrublands (Table 1). Spatially averaged DSI also exhibited a negative correlation with MIT and EMIT in forests (Table 1). These findings indicated that climatic warming may result in drying at ecosystem scale, which was in line with previous studies [28-30].

Spatially averaged DSI exhibited a negative correlation with SP and SH in alpine meadows, alpine steppes, croplands, forests and shrublands (Table 1), which implied that the increase of sunshine may cause drying at ecosystem scale.

Generally, correlations between DSI and climatic factors varied among the 69 stations (Figure S4). Three stations were mainly pre-dominated by wind speed, 26 stations were mainly pre-dominated by temperature factors; 11 stations were mainly pre-dominated by precipitation factors; 15 stations were mainly pre-dominated by air humidity factors; and 14 stations were mainly pre-dominated by sunshine factors. Most stations exhibited positive correlations between DSI and water availability, and negative correlations between DSI and sunshine conditions. Most stations also exhibited negative correlations of DSI with MAT and EMAT. More than half stations exhibited negative correlations of DSI with Ta, MIT and EMIT. These findings indicated that DSI can mirror water availability at station scale. Moreover, both climatic warming and the increase of sunshine resulted in drying at most stations. This finding was in line with previous studies conducted on the Tibetan Plateau [31-33]. For example, Yu et al. [34] demonstrated that experimental warming increased soil temperature by approximately 1−1.4 °C, but decreased soil moisture by approximately 4% in an alpine meadow in Northern Tibet.

## 4. Conclusions

The main conclusions are as follows: (1) different vegetation types exhibited different correlations between DSI and climatic factors, which may be attributed to different changes and background values in climatic factors among vegetation types; and (2) the degree of correlation between DSI and climatic change was stronger in colder, drier, higher wind speed and sunshine environments.

## Figure legend

**Figure 1.** Linear trends for drought severe index (DSI) from 2000 to 2011 over the entire Tibetan Plateau

**Figure 2.** Drought severity index trends from 2000 to 2011 on the Tibetan Plateau; (a) significance test; and (b) regression slope.

**Figure 3.** Relationships between several factors: (a) linear trend of annual drought severe index (Slope_DSI) and linear trend of annual maximum temperature (Slope_MAT); (b) Slope_DSI and linear trend of annual extremely maximum temperature (Slope_EMAT); (c) Slope_DSI and linear trend of annual extremely minimum temperature (Slope_EMIT); (d) Slope_DSI and linear trend of annual precipitation (Slope_TP); (e) Slope_DSI and linear trend of annual sunshine percentage (Slope_SP); and (f) Slope_DSI and linear trend of annual sunshine hours (Slope_SH).

## Acknowledgments

We thank the Data Sharing Infrastructure of Earth System Science in China and the China Meteorological Data Sharing Service System for providing vegetation type maps and climatic data, respectively. This work was supported by the National Key Research Projects of China [Nos. 2017YFA0604801; 2016YFC0502006], National Natural Science Foundation of China [Nos. 31600432, 41571042, 41761008], the Science and Technology Plan Projects of Tibet Autonomous Region (Forage Grass Industry) and Youth Innovation Research Team Project of Key Laboratory of Ecosystem Network Observation and Modeling (LENOM2016Q0002).

## References

1. Dai AG (2011) Drought under global warming: a review. Wiley Interdisciplinary Reviews-Climate Change 2: 45–65.

2. Mu QZ, Zhao MS, Kimball JS, McDowell NG, Running SW (2013) A remotely sensed global terrestrial drought severity index. Bulletin of the American Meteorological Society 94: 83–98.

3. Trenberth KE, Dai AG, van der Schrier G, Jones PD, Barichivich J, et al. (2014) Global warming and changes in drought. Nature Climate Change 4: 17–22.

4. Fu G, Zhang HR, Sun W (2019) Response of plant production to growing/non-growing season asymmetric warming in an alpine meadow of the Northern Tibetan Plateau. Science of the Total Environment 650: 2666–2673.

5. Wang M, Zhou CP, Wu L, Xu XL, Ou YH (2013) Wet-drought pattern and its relationship with vegetation change in the Qinghai-Tibetan Plateau during 2001-2010. Arid Land Geography 36: 49–56.

6. Fu G, Sun W, Yu CQ, Zhang XZ, Shen ZX, et al. (2015) Clipping alters the response of biomass production to experimental warming: a case study in an alpine meadow on the Tibetan Plateau, China. Journal of Mountain Science 12: 935–942.

7. Fu G, Shen ZX, Zhang XZ (2018) Increased precipitation has stronger effects on plant production of an alpine meadow than does experimental warming in the Northern Tibetan Plateau. Agricultural and Forest Meteorology 249: 11–21.

8. Dai AG (2013) Increasing drought under global warming in observations and models. Nature Climate Change 3: 52–58.

9. Sheffield J, Wood EF, Roderick ML (2012) Little change in global drought over the past 60 years. Nature 491: 435-+.

10. Yu CQ, Wang JW, Shen ZX, Fu G (2019) Effects of experimental warming and increased precipitation on soil respiration in an alpine meadow in the Northern Tibetan Plateau. Science of the Total Environment 647: 1490–1497.

11. Fu G, Shen ZX, Sun W, Zhong ZM, Zhang XZ, et al. (2015) A meta-analysis of the effects of experimental warming on plant physiology and growth on the Tibetan Plateau. Journal of Plant Growth Regulation 34: 57–65.

12. IPCC (2013) Summary for Policymakers. In: Climate Change 2013: The Physical Science Basis. Contribution of Working Group I to the Fifth Assessment Report of the Intergovernmental Panel on Climate Change [Stocker, T.F., D. Qin, G.-K. Plattner, M. Tignor, S.K. Allen, J. Boschung, A. Nauels, Y. Xia, V. Bex and P.M. Midgley (eds.)]. Cambridge University Press, Cambridge, United Kingdom and New York, NY, USA.

13. Wu JS, Fu G (2018) Modelling aboveground biomass using MODIS FPAR/LAI data in alpine grasslands of the Northern Tibetan Plateau. Remote Sensing Letters 9: 150–159.

14. Seager R, Naik N, Vecchi GA (2010) Thermodynamic and Dynamic Mechanisms for Large-Scale Changes in the Hydrological Cycle in Response to Global Warming. Journal of Climate 23: 4651–4668.

15. Zhong ZM, Shen ZX, Fu G (2016) Response of soil respiration to experimental warming in a highland barley of the Tibet SpringerPlus 5: doi: 10.1186/s40064-40016-41761-40060.

16. Shen ZX, Wang JW, Sun W, Li SW, Fu G, et al. (2016) The soil drying along the increase of warming mask the relation between temperature and soil respiration in an alpine meadow of Northern Tibet. Polish Journal of Ecology 64: 125–129.

17. Wang L, Chen W (2014) A CMIP5 multimodel projection of future temperature, precipitation, and climatological drought in China. International Journal of Climatology 34: 2059–2078.

18. Shen ZX, Fu G, Yu CQ, Sun W, Zhang XZ (2014) Relationship between the growing season maximum enhanced vegetation index and climatic factors on the Tibetan Plateau. Remote Sensing 6: 6765–6789.

19. Fu G, Li SW, Sun W, Shen ZX (2016) Relationships between vegetation carbon use efficiency and climatic factors on the Tibetan Plateau. Canadian Journal of Remote Sensing 42: 16–26.

20. Wang SH, Sun W, Li SW, Shen ZX, Fu G (2015) Interannual variation of the growing season maximum normalized difference vegetation index, MNDVI, and its relationship with climatic factors on the Tibetan Plateau. Polish Journal of Ecology 63: 424–439.

21. Liu DK, Wang JB, Qi SH (2014) Analysis on dry trend based on moisture index in Qinghai province in the recent 35 years. Research of Soil and Water Conservation 21: 246–250.

22. Yang XH, Zhuo G, Luo B (2014) Drought monitoring in the Tibetan Plateau based on MODIS dataset. Journal of Desert Research 34: 527–534.

23. Fu G, Wu JS (2017) Validation of MODIS Collection 6 FPAR/LAI in the alpine grassland of the Northern Tibetan Plateau. Remote Sensing Letters 8: 831–838.

24. Zhang XZ, Shen ZX, Fu G (2015) A meta-analysis of the effects of experimental warming on soil carbon and nitrogen dynamics on the Tibetan Plateau. Applied Soil Ecology 87: 32–38.

25. Fu G, Shen ZX (2017) Effects of enhanced UV-B radiation on plant physiology and growth on the Tibetan Plateau: a meta-analysis. Acta Physiologiae Plantarum 39: doi: 10.1007/s11738-11017-12387-11738.

26. Wang SH, Sun W, Li SW, Shen ZX, Fu G (2015) Interannual variation of the growing season maximum normalized difference vegetation index, MNDVI, and its relationship with climatic factors on the Tibetan Plateau. Polish Journal of Ecology 63: 291–306.

27. Zhang L, Guo HD, Ji L, Lei LP, Wang CZ, et al. (2013) Vegetation greenness trend (2000 to 2009) and the climate controls in the Qinghai-Tibetan Plateau. Journal of Applied Remote Sensing 7: doi:10.1117/1111.jrs.1117.073572.

28. Yu CQ, Han FS, Fu G (2019) Effects of 7 years experimental warming on soil bacterial and fungal community structure in the Northern Tibet alpine meadow at three elevations. Science of the Total Environment 655: 814–822.

29. Fu G, Shen ZX, Zhang XZ, Zhou YT (2012) Response of soil microbial biomass to short-term experimental warming in alpine meadow on the Tibetan Plateau. Applied Soil Ecology 61: 158–160.

30. Fu G, Shen ZX (2017) Clipping has stronger effects on plant production than does warming in three alpine meadow sites on the Northern Tibetan Plateau. Scientific Reports 7: doi: 10.1038/s41598-41017-16645-41592.

31. Fu G, Shen ZX (2016) Environmental humidity regulates effects of experimental warming on vegetation index and biomass production in an alpine meadow of the Northern Tibet. PLoS ONE 11: 10.1371/journal.pone.0165643.

32. Shen ZX, Li YL, Fu G (2015) Response of soil respiration to short-term experimental warming and precipitation pulses over the growing season in an alpine meadow on the Northern Tibet. Applied Soil Ecology 90: 35–40.

33. Fu G, Zhang XZ, Zhang YJ, Shi PL, Li YL, et al. (2013) Experimental warming does not enhance gross primary production and above-ground biomass in the alpine meadow of Tibet. Journal of Applied Remote Sensing 7: 10.1117/1111.jrs.1117.073505.

34. Yu CQ, Shen ZX, Zhang XZ, Sun W, Fu G (2014) Response of soil C and N, dissolved organic C and N, and inorganic N to short-term experimental warming in an Alpine meadow on the Tibetan Plateau. Scientific World Journal 2014: 10.1155/2014/152576.

